# “Comparative analysis of red blood cell under real and simulated microgravity conditions”

**DOI:** 10.1101/2024.12.26.630384

**Authors:** Sarika Hinge, Jyotsana Dixit, Pandit Vidyasagar, Gauri Kulkarni

## Abstract

Red blood cell is an important role model of microcirculation, gas exchange, drug carrier, and related diseases. There has been an increasing interest in understanding the biochemical and structural changes in Red blood cells (RBC) due to simulated microgravity conditions. In the present work, isolated human red blood cells were exposed to simulated microgravity (SMG) conditions using 2D clinostat. Comparative analysis of normal RBC and RBC in simulated microgravity conditions was performed using UV-visible spectroscopy, FTIR spectroscopy, and single-beam optical tweezer. The change in position of the Soret band absorption peak is observed after the exposure to simulated microgravity. However, biochemical functional groups of RBCs were studied using FTIR spectroscopy with a change in % Transmittance. Power spectrum density (PSD) variations in optically trapped control and SMG exposed single RBC are obtained from position signals acquired by Thorlabs’s OTKBFM-CAL data acquisition software. Microgravity (SMG)-induced stress is responsible for a reduction in haematocrit, haemolysis, change in the morphology, and rigidity of red blood cells. These investigation of the physicochemical properties of RBC in simulated microgravity is useful for improving the health of astronauts during space missions.

## Introduction

Response of red blood cells to external physical factors viz. gravity [1], radiation [2-3], vibration [4], electromagnetic field [5], osmotic pressure [6], and temperature [7] plays an environmental parameters to study basic processes in life sciences. However, microgravity differs from normal earth gravity [8]. During space exploration, the health and safety of astronauts is important. Hence, it is necessary to understand the cell growth, cell cycle, differentiation, proliferation, morphogenesis, interaction mechanism, etc. in simulated microgravity/ altered gravity conditions whether biological system experienced near weightlessness within the range of 10^-3^ to 10^-6^ g [9]. Simulated microgravity is used to investigate the effect of microgravity on biological cells, plants, seeds, bacteria, bones, tissues, macrophages, etc [10-15]. Cells subjected to microgravity conditions exhibits the changes in cytoskeletal organization, and mechanotransduction [16-17]. The alteration or modification of biological cells depends on exposure time, rotational speed, angular frequency, and physical parameters like temperature, humidity, % CO_2_, etc [18].

To perform the space experiment on earth, microgravity is simulated using 2 D clinostat, and random positioning machine (RPM) [19-20]. SMG induces stress on biological cells [21]; thus, RBC is used as a model system for pathophysiological study. RBC is composed of haemoglobin, water, membrane protein, and spectrin network composed of cytoplasmic skeleton [22]. RBC experiences shear stress during microcirculation under normal gravity condition. The effect of microgravity on blood cells has the potential to understand the blood cell biology and manned space flight. On earth, blood flows towards the leg in upright position; however, in space blood flows from leg to brain in a reverse cyclic manner in microgravity which affects the whole-body function, fluid dynamics, haemorheological properties, and as a result cardiovascular disorder, intracranial hypertension, and blood vessel constriction [23]. In order to explore the pathophysiology and biochemistry in cell development of living things, a variety of approaches, including numerical simulations, microfluidics, organ on a chip, and organoids, were utilized to examine the implications, assessment, and evaluation of microgravity on animals and plants [24-26]. Furthermore, several biochemical tests for amylase activity, aging, and lipid peroxidation research are conducted, along with optical microscopy, atomic force microscopy (AFM), fluorescence microscopy, confocal microscopy, flow cytometry, and spectroscopic techniques [27]. In the present work, UV-visible spectroscopy, FTIR spectroscopy, and optical tweezer were used for studying the effect of Simulated Microgravity on Red Blood Cells. These findings enable a better understanding of red blood cell and its oxygenation state in a microgravity environment to improve health outcomes for our pilots and astronauts.

## Material and methods

Requirement: Human Adult blood, EDTA tubes phosphate buffer saline 127 mM NaCl, 2.7 mM KCl, 8.1 mM Na_2_HPO_4_, 1.5 mM KH_2_PO_4_, Deionized water, pH 7.4).

### Collection and Isolation of RBC from whole blood

Whole blood from healthy adults (Hb:14) was drawn from a specialized medical expert. The patient provided written, informed consent in before performing experiments. This study was conducted with ethical permission and the Declaration of Helsinki. Blood was collected from the EDTA tube and stored at 4 °C before used. RBCs were isolated by centrifugation method at 3000 RPM for 10 min. Due to the high-density gradient, RBCs were settled down at the bottom of the tube. Isolated RBCs were washed thrice with phosphate buffer saline to avoid the debris and maintain its physiological condition. RBC suspension (2%) was used to experiment. To understand the mechanism of RBC in simulated microgravity, experiments were performed in a 2 D clinostat (Department of Physics, Savitribai Phule Pune University, Pune, India) for 12 hr, and examined using UV-visible spectroscopy, FTIR spectroscopy, and Optical Tweezer.

### Experimental Set up

For simulation of microgravity, 2-D clinostat instrument was used. It consists of stepper motor (12V) embedded with sample chamber for microgravity exposure. One fixed sample chamber was assembled for normal Gravity (1g). Whole set up kept in an environment chamber with controlled temperature and humidity. Suspension of RBC (2 %) were used for microgravity exposure as shown in Figure (1).

**Figure 1:**
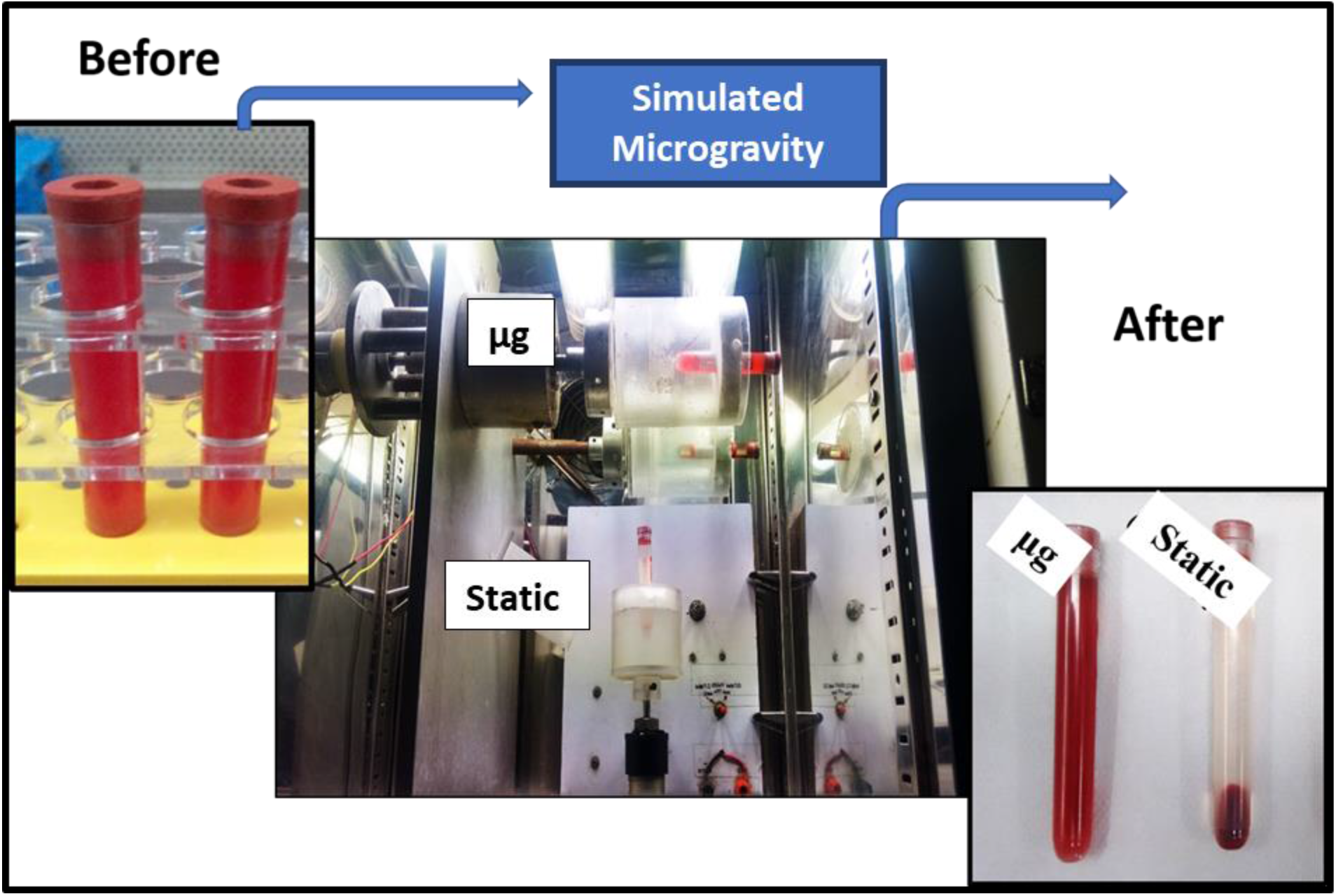
Simulated Microgravity Set up to expose red blood cells (Before and after simulated RBC))

### UV-visible absorption spectroscopy

UV-visible absorption spectra of RBC suspension were recorded to confirm the changes in Hb group and its associated oxygen binding sites. Figure (2) depicts the comparison spectra of RBC at static condition and simulated microgravity (SMG) exposed RBC. Absorption spectra of static and SMG exposed RBC showed the characteristics peaks at 280 nm, 341 nm, 418 nm, 541 nm, 579 nm. The peaks at 280 nm corresponds to protein, 341 nm corresponds to tyrosine, 418 nm represents the haem group and oxygen carrying capacity shown by Q1, Q2 i.e. at 541 nm and 579 nm [28]. From figure 2 (a-d) it is seen that, microgravity affects the overall RBC function with decrease in absorbance.

**Figure 2:**
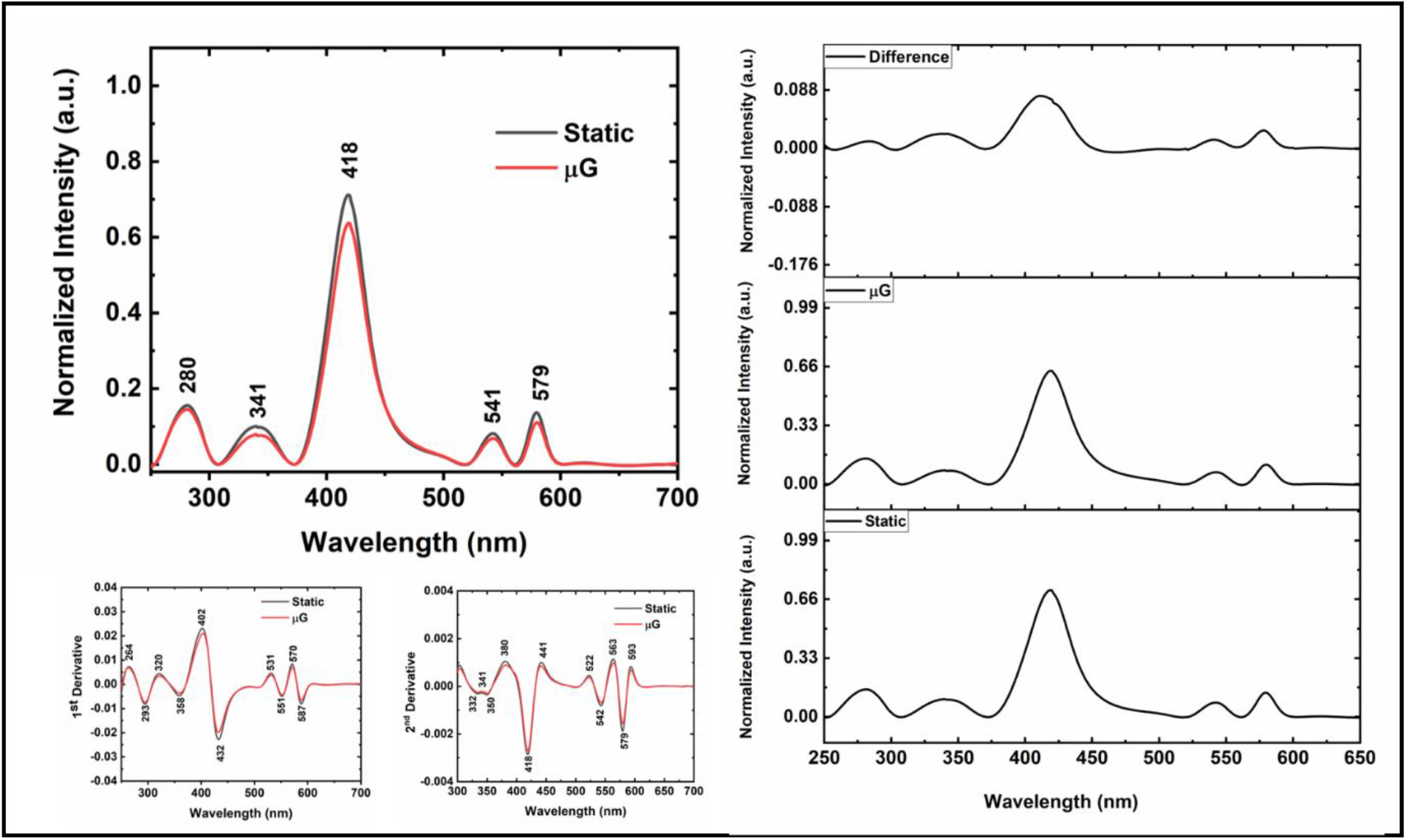
(a) UV-visible absorption spectroscopy of Microgravity simulated red blood cells (b) difference (c) 1^st^ derivative (d)2^nd^ derivative

### FTIR spectroscopy

The FTIR spectra of RBC in static condition and SMG exposed RBC were collected within the range of 500 cm^-1^ to 4000 cm^-1^ using a Savitzky-Golay method. The functional group assignment of RBC was done according to previous research paper [29]. RBC suspension (2%) were exposed to microgravity under simulated condition and immediately obtained and compared the FTIR spectra with static.

FTIR spectra of RBC showed the peaks at 1271 cm^-1^, 1415 cm^-1^, 1544 cm^-1^, 1640 cm^-1^, 2067 cm^-1^, and 3423 cm^-1^ under normal gravity (static g) and SMG (µg) as shown in Figure 3 (a). The first derivative of transmission spectra of RBC revealed the peak position at 1235 cm^-1^, 1338 cm^-1^, 1375 cm^-1^,1391 cm^-1^,1404 cm^-1^, 1425 cm^-1^, 1462 cm^-1^, 1513 cm^-1^, 1534 cm^-1^, 1551 cm^-1^, 1601 cm^-1^ and 1689 cm^-1^. Figure 3(b) depicts the reduction in transmission of simulated microgravity exposed RBC. These reductions are due to changes occur at polypeptide chain, amide I and amide II. Degradation of RBC membrane is found in second derivatives of FTIR spectra. FTIR spectra recorded in Figure 3(c-d) shows enhancement in intensity from 1620 cm^-1^ to 1660 cm^-1^ in Amide I with shift in wavenumber which indicates that change in the secondary structure viz. α-helix, β-turns, and intermolecular β-sheets. As a result, modification in proteins present in the RBC membrane which are associated with spectrin and actin. Shift in wavenumber and reduction in intensity of SMG exposed RBC is observed within the spectral range of 1675 cm^-1^ to 1700cm^-1^ and 2800 cm^-1^ to 3000 cm^-1^.

**Figure 3:**
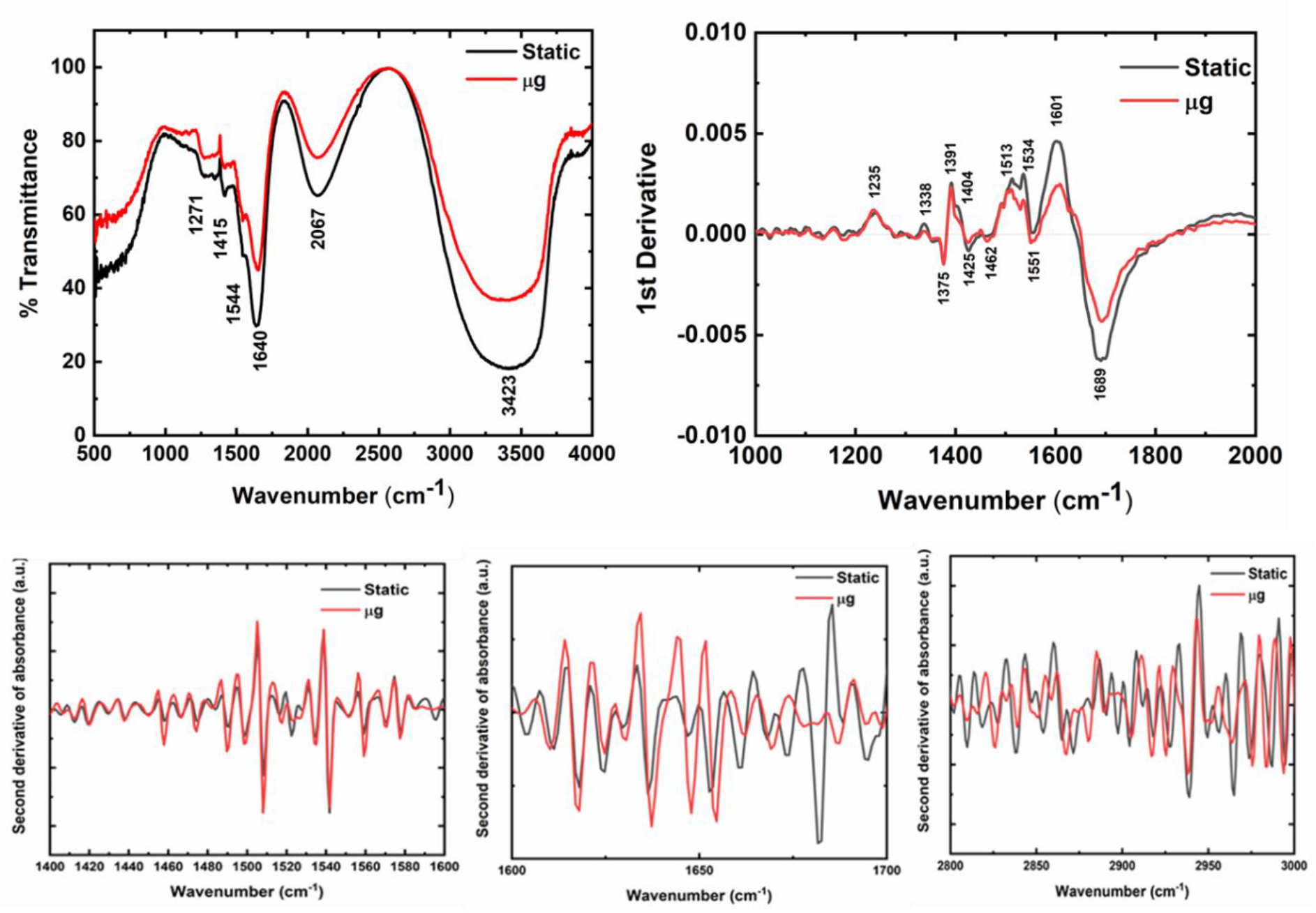
(a) FTIR of simulated microgravity exposed RBC and (b) its 1^st^ derivative

### Optical Trapping

Thorlab’s modular optical tweezer is used to hold a RBC with wavelength 975 nm and power 100 mW.

Figure 4 shows the PSD spectra of optically trapped simulated microgravity exposed single RBC. Decrease in the power spectrum of red blood cell was observed in simulated microgravity-exposed RBC due to change in RBC membrane properties.

**Figure 4:**
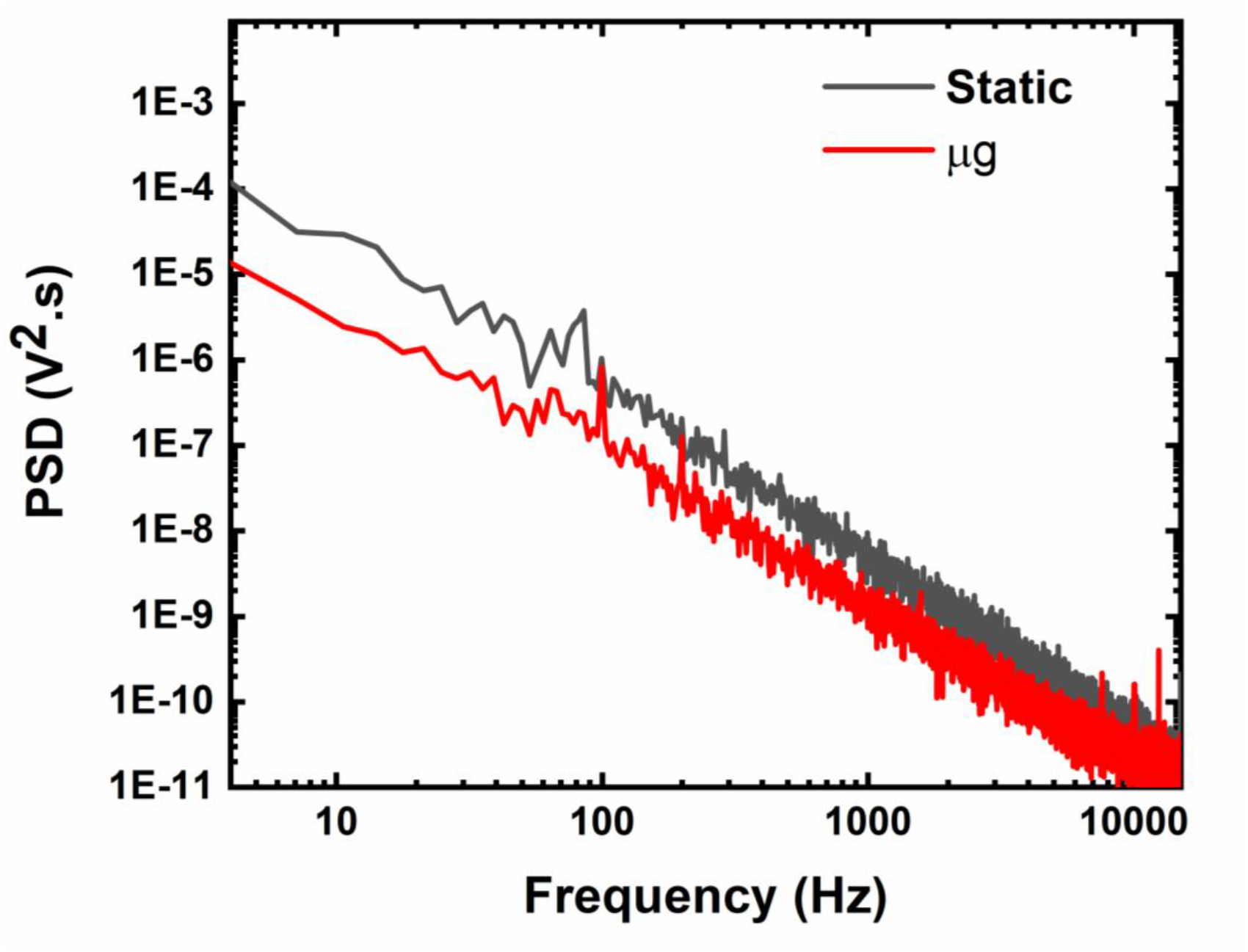
Optical trapping of microgravity simulated Red blood cell

## Discussion

The Soret band present at 418 nm attributed to haem group which is enclosed by porphyrin ring and polypeptide chain. In case of Hb structure, central position of porphyrin ring is occupied by iron group with six co-ordinated sites. Along with oxygen binding sites, histidine and two alpha and two beta polypeptide chains. No shift in spectral position of RBC were observed in SMG exposed RBC. Change in spectral intensity showed the decrease in Hb which suggest the rupture of RBC as a results haemolysis occur in µg treated RBCs. Studies have shown that exposure to microgravity can lead to a reduction in red blood cell mass. This means that astronauts may experience a decrease in the total number of red blood cells in their bodies. ^59^Fe, ^51^Cr radio-label RBC were used to study changes in haemoglobin parameter associated with a decrease in the size and quantity of red blood cells produced during spaceflight [30]. This reduction of hematocrit can affect the oxygen-carrying capacity of the blood. The decrease in intensity at Q band i.e. at 542 nm and 576 nm represent the signature peak of oxygenation state.

In normal static condition, RBCs were settled down at the bottom; however, in simulated µg condition, RBCs present throughout the PBS suspension i.e. no sedimentation. Thus, microenvironment of static and SMG exposed RBC is differed. The effect of microgravity on RBC dynamics influences the size, shape, volume, fluidity, osmotic pressure, membrane permeability, adhesion, and aggregation. It also affects the binding sites of haemoglobin function of oxygen transport, cell transduction, and homeostasis. NASA held a project on the microgravity effect of blood, results showed that change in morphological behavior in red blood cells, plasma volume, bone marrow, and erythropoietic. No sedimentation occurs in cell is observed during BIOMICS project experiment (MASER I and II). Microgravity causes atrophy of the muscular system, dysfunction of the immune system, electrolyte imbalance, cardiovascular anomalies, reduction in bone mass, and osteopenia [31]. Microgravity induces oxidative stress on RBCs which leads to an increase of haematocrit and Hb concentration, causing the destruction of immature RBCs in the bone marrow and its oxygen carrying capacity.

Lift force and hydrodynamic interaction play an important role in the shear-induced fluid rheology of blood, and modification of mechanical properties [32-33]. A decrease in hydrostatic pressure alerts the mechanical behavior of RBC as a result of deformation which causes an insufficient supply of oxygen to tissues. Cells may experience altered deformation and changes in shape due to reduced gravitational forces. This can impact their ability to maintain a normal morphology and interact with neighboring cells and surfaces. Microgravity-exposed cells experienced stress which caused the rupture of the cell wall or modification in cell membrane. Thus, a higher rate of haemolysis and ageing was observed as compared to the normal gravity condition [34]. The first and second derivative of FTIR spectra of normal and simulated microgravity exposed RBC showed variation in secondary structure, alteration in proteins, phospholipids, and unsaturated fatty acid. Decrease in the cell membrane roughness is related to aging in microgravity. In current study, SMG induced changes in RBC membrane is observed PSD spectra which is obtained using Optical Tweezer technique.

## Conclusion

In this work, we investigate the spectroscopic analysis of microgravity-exposed RBC. UV-visible spectroscopy analysis demonstrates that simulated microgravity-exposed RBC showed changes in the haem group or Soret band and its oxygenation state. First derivative of FTIR spectroscopy showed decrease in % transmittance. Simulated microgravity affects the power spectrum activity of the Red blood cell which is an indication of an alteration in the cell membrane stiffness. Microgravity can lead to changes in the shape, deformability, and function of red blood cells. Future studies will provide further evidence of the microgravity approach in nanomedicine.

## Authors Contributions

Conceptualization, S.B.; methodology, S.B. and J.D.; writing-original draft: S.B.; writing - review and editing, S.B., J.D., P.V., and G.R.; Supervision: G.R; All authors have read and agreed to the published version of the manuscript.

## Acknowledgment

Author thankful to the School of Basic Medical Sciences, Department of Physics, Savitribai Phule Pune University, Pune.

## Ethics Declaration

This study is approved by the Institutional Ethical Committee (Ref no. SPPU/IES/2021/129). Written consents were taken from individual subjects.

## Data availability statement

All relevant data are within the manuscript.

